# RubiSpec-MS: Determination of Rubisco CO_2_/O_2_ Specificity Factor using Liquid Chromatography-Mass Spectrometry

**DOI:** 10.64898/2026.06.14.731917

**Authors:** April Wu, Yunlong Zhao, Julie L. McDonald, Mohanraja Kumar, Robert H. Wilson, Matthew D. Shoulders

## Abstract

- The specificity factor(*S*_C/O_) of ribulose-1,5-bisphosphate carboxylase/oxygenase (Rubisco) is a key kinetic parameter quantifying the ratio of carboxylation to oxygenation reactions. Current methods to measure *S*_C/O_ are expensive, labor-intensive, require specialized expertise, and/or rely on radiolabeled compounds, hindering Rubisco research.
- We present RubiSpec-MS (Rubisco Specificity by LC-MS/MS), a method to determine the *S*_C/O_ of purified Rubiscos using liquid chromatography-coupled mass spectrometry (LC-MS/MS) for separation and sensitive detection of non-radioactive reaction products.
- We benchmark RubiSpec-MS using multiple model Rubisco enzymes. We find that the assay accurately reproduces corresponding literature values for each enzyme and performs similarly in a side-by-side comparison using an established radiometric method.
- RubiSpec-MS provides a faster, more accessible, and highly parallelizable method to measure Rubisco *S*_C/O_ using equipment common in academic core facilities and industrial laboratories.

## INTRODUCTION

Ribulose-1,5-bisphosphate carboxylase/oxygenase (Rubisco) is the central enzyme in the Calvin–Benson–Bassham (CBB) cycle, which is responsible for nearly all biological carbon fixation on Earth (Prywes *et al*., 2023). While carbon fixation is the main function of Rubisco, the enzyme can also catalyze oxygenation, owing to a failure to properly discriminate between CO_2_ and O_2_. Both carboxylation and oxygenation proceed via reaction with Rubisco’s substrate ribulose-1,5-bisphosphate (RuBP). Carboxylation generates two molecules of 3-phosphoglycerate (3PGA), which are used to regenerate RuBP and produce metabolite sugars, such as glucose. Oxygenation generates one molecule of 3PGA and one molecule of 2-phosphoglycolate (2PG). In plants, 2PG must then be converted to 3PGA through photorespiration, a process that releases CO_2_ and consumes energy (**Fig. 1a**) (Walker *et al*., 2016).

**Figure 1.**
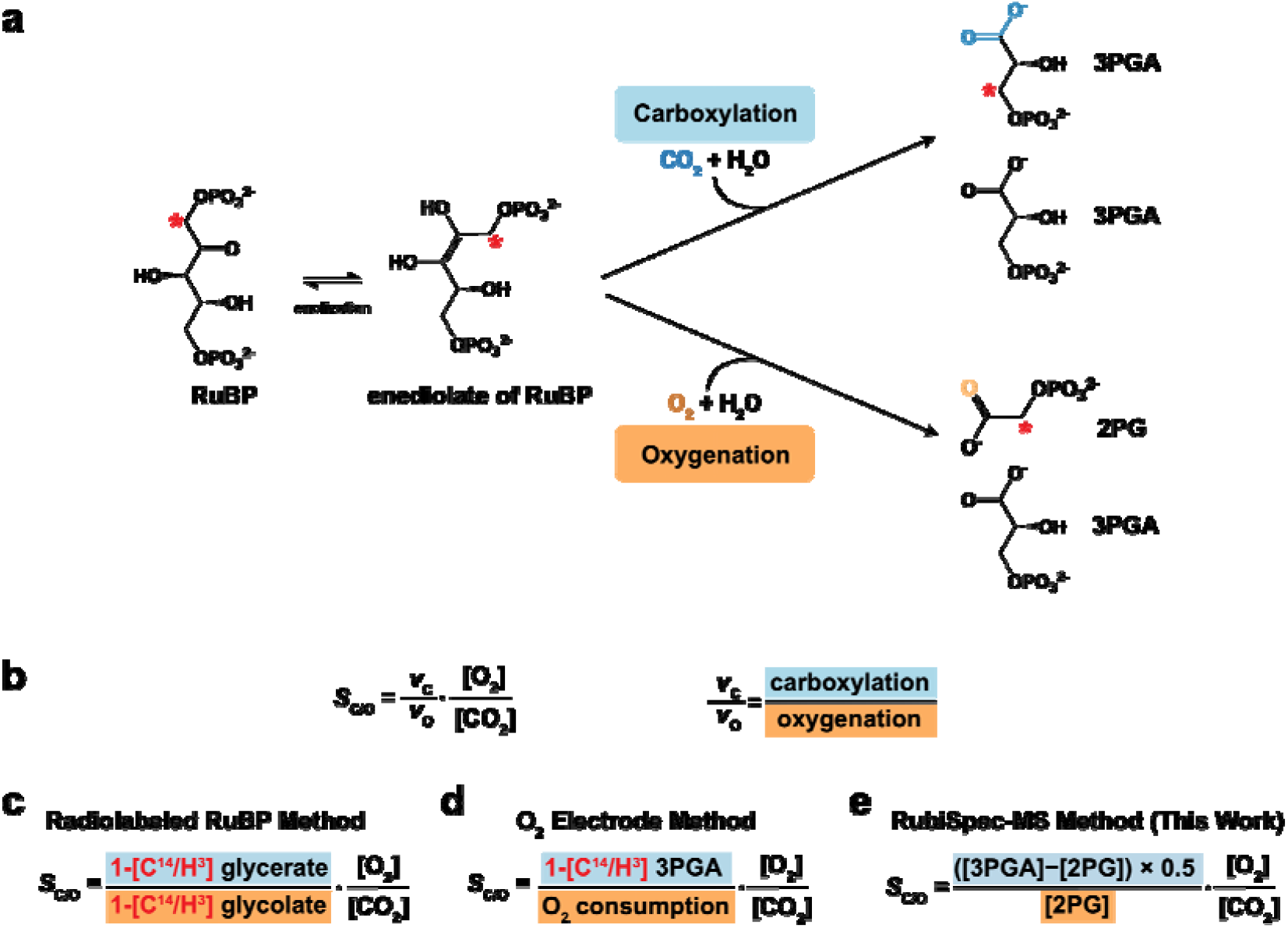
Summary of Rubisco carboxylation and oxygenation reactions and methods to measure *S*_C/O_. **(a)** Rubisco first catalyzes the enolization of RuBP before the addition of gaseous sub-strate. Carboxylation generates two molecules of 3PGA and oxygenation generates one moleculeed by red asterisks. **(b)** Rubisco specificity factor (*S*_C/O_) is calculated as the ratio of carboxyla-of 2PG and one molecule of 3PGA. ^14^C positions for the radiolabeled RuBP method are indicatsite. **(c)** Calculation of *S*_C/O_ from the Radiolabeled RuBP method relies on quantification of radition to oxygenation rate multiplied by the concentration of O_2_ to CO_2_ gas at the Rubisco active olabeled glycerate and glycolate. **(d)** Calculation of *S*_C/O_ from the O_2_ Electrode method relies on quantification of radiolabeled glycerate and O_2_ consumed during the reaction. **(e)** Calculation of *S*_C/O_ from the RubiSpec-MS method relies on quantification of unlabeled 3GPA and 2PG. Abbreviations: RuBP (ribulose-1,5-bisphosphate), 2PG (2-phosphoglycolate), 3PGA (3-phosphoglycerate).

Rubisco’s slow carboxylation rate and competition by O_2_ limit photosynthetic efficiency, making it the subject of extensive biochemical research and genetic engineering efforts to enhance plant productivity (Long *et al*., 2006; Whitney *et al*., 2011). Such efforts depend on accurate and precise determination of the kinetic parameters that describe the carboxylation and oxygenation reactions. Accordingly, Rubisco kinetic characterization has focused on three key quantities: the maximum carboxylation rate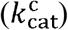, the Michaelis–Menten constant for CO_2_ (*K*_C_), and the specificity factor (*S*_C/O_), which describes the relative gas substrate specificity of Rubisco for CO_2_ over O_2_. The direct Michaelis-Menten parameters for oxygenation 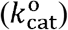 and *K*_O_ are also relevant, but they are reported less often. In particular, 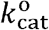 is difficult to determine directly because Rubisco requires the presence of CO_2_ to both form and maintain the active site throughout the assay. Moreover, there is no convenient radioisotope for 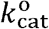 measurement. Instead, 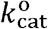 is typically derived indirectly from other kinetic constants and rearrangement of the equation *S*_C/O_ (**Fig. 1b**). Consequently, in most cases, determination of 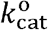 and *K*_O_ are dependent on reliable methods to measure *S*_C/O_.

Rubisco’s key kinetic parameters are critical for studying the enzyme’s catalytic mechanism (Wilson & Whitney, 2017; Flamholz *et al*., 2019), quantifying differences in Rubisco behavior across natural systems (Galmés *et al*., 2005; Orr *et al*., 2016); Young *et al*. (2016); (Iñiguez *et al*., 2020; Aguiló-Nicolau *et al*., 2024), and enabling engineering efforts to optimize the enzyme (Wilson *et al*., 2018; Zhou & Whitney, 2019; Chen *et al*., 2022; Gionfriddo *et al*., 2025; McDonald *et al*., 2025). Accurate knowledge of Rubisco kinetic parameters is also essential for modeling photosynthetic efficiency (Farquhar *et al*., 1980; Brooks & Farquhar, 1985; Bernacchi *et al*., 2001; Busch, 2020), improving crop yields through genetic engineering, and informing Earth system models that predict the impacts of rising atmospheric CO_2_ and temperature on gross primary production (Bowes, 1991; Rogers, 2014) and climate change (Galmés *et al*., 2013; Prywes *et al*., 2023). More recently, some of these same kinetic parameters have also been used as training data to inform machine learning approaches to Rubisco discovery and engineering (Iqbal *et al*., 2023; Prywes *et al*., 2023). Clearly, accessible Rubisco kinetic assays are paramount to support climate modeling and agricultural engineering efforts.

Several assays have been developed to quantify various aspects of Rubisco kinetics. 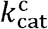can be determined non-radiometrically via a spectrophotometric NADH-coupled enzyme assay that is compatible with a wide range of purified Rubiscos (Kubien *et al*., 2011; Sharwood *et al*., 2016; Young *et al*., 2016; Davidi *et al*., 2020).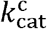 and *K*_C_ can be determined simultaneously by fitting the substrate concentration versus carboxylation rate curve at varying CO_2_ concentrations, most often by measuring ^14^C incorporation into 3PGA in the absence of O_2_ (Iñiguez *et al*., 2021). The use of radioisotopes permits these measurements using even crude preparations of Rubisco from cellular lysates.*K*_O_ can be determined via the same ^14^C-based assay, by measuring the inhibition of carboxylase activity at varying O_2_ concentrations.

Amongst Rubisco kinetic parameters, determination of *S*_C/O_ is particularly challenging because it requires precision control over the atmosphere of the reaction and specialized equipment and reagents that are not commercially available and typically must be synthesized inhouse. Thus, *S*_C/O_ measurements are most often performed separately from other kinetic assessments of activity. The most common methods are radiometric. The first, hereafter referred to as the ‘Radiolabeled RuBP Method’, uses RuBP radiolabeled at the C-1 position ([1-^14^C/^3^H]-RuBP) such that gas addition at C-2 will always lead to the production of ^14^C-labeled 3-phosphoglycerate (3PGA) from CO_2_ and ^14^C-labeled 2-phosphoglycolate (2PG) from O_2_ (**Fig. 1a**) (Kane *et al*., 1994). Given known mole fractions of CO_2_ and O_2_ in the reaction, the ratio of CO_2_ to O_2_ fixed can be determined via separation of the products followed by sensitive radiometric quantification of radiolabeled glycerate and glycolate, respectively (**Fig. 1c**). This method is highly accurate and can be performed using minimal amounts of purified Rubisco enzyme. Unfortunately, the need for in-house synthesized, radiolabeled RuBP poses a significant barrier to those not experienced with synthesis or use of radioactive materials, and the needs for operation of precision gas mixing devices and management of radioactive waste make it further challenging to implement.

The second method (Parry *et al*., 1989), hereafter referred to as the ‘O_2_ Electrode Method’, measures O_2_ fixation using a Clark electrode and CO_2_ fixation via post-reaction scintillation counting of the amount of ^14^C incorporated into [1-^14^C]-3PGA formed from commercially available NaH^14^CO_3_ (**Fig. 1d**). In this approach, *S*_C/O_ of previously uncharacterized Rubisco are normalized to an internal standard, usually a well-characterized Rubisco with a similar expected value (Valegård *et al*., 2018). The O_2_ Electrode Method is relatively simpler and quicker than the Radiolabeled RuBP Method but is still highly specialized. The use of aqueous NaH^14^CO_3_ represents a significant safety hazard, as it can generate gaseous ^14^CO_2_ and must be handled under a fume hood.

Given the labor-intensive nature and hazards of these methods, several groups have sought alternatives. Accordingly, Rubisco specificity assays that rely on nuclear magnetic resonance (Wang *et al*., 1998), gas chromatography mass spectrometry (Whitney & Andrews, 1998), membrane inlet mass spectrometry (Cousins *et al*., 2010), and isothermal titration calorimetry (Frank & Müh, 2025) have all been introduced. These methods still face various challenges that limit generalizability, owing to requirements for specialized equipment, low-throughput, high error, or some combination of these and other issues.

Hence, Rubisco *S*_C/O_ determination remains far from routine, posing a significant barrier to entry for researchers potentially interested in the Rubisco field, and even limiting many research groups deeply committed to studying and engineering Rubisco. Currently, only a few laboratories worldwide produce consistent, reproducible measurements of *S*_C/O_.

Here, we introduce RubiSpec-MS (Rubisco Specificity by LC-MS/MS) as a straightforward and reliable approach to measure Rubisco *S*_C/O_ using liquid chromatography-coupled mass-spectrometry (LC-MS/MS). The protocol relies on hydrophilic interaction chromatography (HILIC) and tandem mass spectrometry to separate, detect, and quantify the carboxylation and oxygenation reaction products of Rubisco, from which *S*_C/O_ can be calculated (**Fig. 1e**). Here, we describe in detail the RubiSpec-MS experimental procedure, reagents, and data analysis to determine Rubisco *S*_C/O_ for multiple model Rubisco enzymes. We exemplify the reliability of RubiSpec-MS by comparing results from samples analyzed side-by-side with the gold standard Radiolabeled RuBP Method as a direct comparison. The results demonstrate that RubiSpec-MS delivers *S*_C/O_ measurements consistent with the Radiolabeled RuBP Method, as well as with previous *S*_C/O_ values from the literature. RubiSpec-MS does not rely on radiolabeled components and provides a simpler, more parallelizable method to measure Rubisco *S*_C/O_ without compromising accuracy or precision. RubiSpec-MS should enable broader analysis of natural and engineered Rubiscos by diverse laboratories using widely available equipment, providing a valuable resource for carbon fixation research.

## MATERIALS AND METHODS

### Chemicals and Reagents

Acetic acid (CH□CO_2_H), HPLC-grade acetonitrile (ACN), ammonium hydroxide (NH_4_OH), ammonium bicarbonate ((NH□)HCO□), hydrogen chloride (HCl), magnesium chloride (MgCl_2_), dithiothreitol (DTT), phenylmethylsulfonyl fluoride (PMSF), D-ribose-5-phosphate (R5P), adenosine triphosphate (ATP), creatine phosphate, polyvinylpolypyrrolidone (PVPP), 3PGA, and 2PG were purchased from Sigma-Aldrich (MA, USA). NADH, glucose, and triethanolamine were purchased from J.T. Baker (PA, USA). Magnesium acetate and imidazole were purchased from Alfa Aesar (MA, USA). Sodium chloride (NaCl) and sodium hydroxide (NaOH) were purchased from VWR Life Sciences (PA, USA). Glyceric acid was purchased from TCI America (OR, USA). Glycolic acid was purchased from ThermoFisher (MA, USA).

DNase-I (RNase-free) was purchased from New England Biolabs (MA, USA). Carbonic anhydrase (CA), phosphoriboisomerase (PRI), glucose-6-phosphate dehydrogenase (G6PDH), and creatine phosphokinase (CPK) were purchased from Sigma-Aldrich (MA, USA). ‘Commercial RuBP’ was also purchased from Sigma-Aldrich. RuBP was synthesized as previously described (Pierce *et al*., 1980). Radiolabeled RuBP ([1-^3^H]-RuBP) was also synthesized as previously described, except that D-[2-^3^H] glucose was substituted for D-[2-^14^C] glucose (Kane *et al*., 1994). Synthesized RuBP and [1-^3^H]-RuBP was stored in liquid nitrogen to reduce oxidation of RuBP to inhibitory pentodiulose-P_2_ (Kane *et al*., 1998).

### Purification of enzymes from *Escherichia coli* extracts

Genes encoding phosphoribulokinase (PRK), phosphogluconate dehydrogenase (6PGD), and the Rubisco large subunit from *Rhodospirillum rubrum* (RbcM) were cloned into a pHue vector that installs an N-terminal 6×His-ubiquitin tag (^His6^Ubq). The ^His6^Ubq was cleaved by 6×His-ubiquitin protease USP2 (^His6^USP2) in a two-step immobilized metal affinity chromatography (IMAC) procedure, as described previously, to yield native enzyme (Catanzariti *et al*., 2004). RbcM is a Form II Rubisco composed of two large subunits (L□), whereas *Cereibacter sphaeroides* Rubisco (*Cs*LS), *Synechococcus elongatus* PCC 6301 Rubisco (*Syn*LS), and *Nicotiana tabacum* Rubisco (*Nt*LS) are Form I Rubiscos that assemble as L□S□ complexes.

Genes encoding untagged Rubisco from *C. sphaeroides* (*Cs*LS) were cloned into a pTrc vector.

The plasmid pET16-*Nt*LS*Rca*, containing the gene encoding untagged *Nt*LS and Rubisco activase (Rca), was obtained as a generous gift from Spencer Whitney (ANU). The Rca gene was removed to create pET16-*Nt*LS. For affinity protein purification of Form I Rubisco, a low copy pACYC plasmid expression system was used to coexpress a ^His6^Ubq-tagged Rubisco small subunits (^His6^Ubq-RbcS), as described previously for *Syn*LS (Mueller-Cajar & Whitney, 2008).

Substoichiometric incorporation of ^His6^Ubq-RbcS into the assembled L_8_S_8_ holoenzyme allows for purification of the native enzyme complex following ^His6^Ubq cleavage, as described above. Accordingly, the coding sequences for the small subunits of *Cs*LS, *Nt*LS, and *Syn*LS featuring N-terminal ^His6^Ubq-RbcS fusions were coexpressed alongside the untagged subunits to facilitate their IMAC purification. The following plasmids were also generous gifts from Spencer Whitney (ANU): pTrc-*Syn*LS, containing the gene encoding *Synechococcus elongatus* PCC 6301 Rubisco; pCDF-*Nt*Asmbl, containing the genes that encode the set of chloroplast chaperones required for assembly of functional plant Rubisco in *E. coli*; (Buck *et al*., 2022); pACYC-*Syn*-^His6^Ubq-RbcS, containing the gene encoding the small subunit of *Syn*LS, and pUSP2-^His6^USP2 encoding 6×Histagged ubiquitin protease (USP2). The full list of plasmids is listed in **Supplementary Table 1**.

Plasmids were chemically transformed into BL21* DE3 cells (Invitrogen).

All cultures were grown in LB medium at 37 °C with shaking at 180 rpm, except for cells expressing 6PGD, which were grown in 2YT medium under the same conditions. Cultures were grown until they reached an *OD*_600_ of 0.6–0.8, at which point protein expression was induced with 1 mM IPTG at 23 °C for 16 h. Cultures were centrifuged for 30 min at 4,000 × *g* and 4 °C. Cell pellets were then flash-frozen in liquid N_2_ and stored at −80 °C.

PRK was purified as previously described (Wilson *et al*., 2019). USP2 was also purified as previously described (Catanzariti *et al*., 2004). 6PGD, RbcM, CsLS, SynLS, and NtLS were purified as follows: Stored cell pellets were resuspended in 30 mL ice-cold buffer (50 mM Tris-HCl at pH 7.5, 300 mM NaCl, 10% (v/v) glycerol, 1 mM PMSF, 5 mM DTT, and 1 µL of DNase-I) then lysed on ice using a needle-tipped sonicator (17% amplitude with 10 s on/off cycles; Branson). Lysates were centrifuged for 45 min at 20,000 × *g* and 4 °C, and then the cleared supernatant was passed through a 0.22 µm cellulose acetate filter. The filtrate was loaded on 4-mL of HisPur™ Cobalt Resin (Thermo Fisher Scientific) equilibrated with column buffer (50 mM Tris-HCl at pH 7.5, 300 mM NaCl, 10% (v/v) glycerol). After washing with 40 mL of column buffer, proteins were eluted with 2 mL fractions of column buffer containing 300 mM imidazole. For each fraction, 2 µL of the eluted protein was mixed with 50 µL of Pierce™ Bradford Protein Assay Kit (Thermo Fisher), and the protein-positive fractions were pooled. For each 3 mL of eluent, 100 µl of 80% (v/v) glycerol, 10 µl of 1.5 M 2-mercaptoethanol, and 50 µl of USP2 (stock of 2 mg/mL) were added and the solution was incubated at 30 °C in a water bath for 30 min. The cleaved protein was passed over regenerated HisPur™ Cobalt Resin and eluted with 6 mL of dialysis buffer (50 mM Tris-HCl at pH 7.5, 50 mM NaCl, 10% (v/v) glycerol) then dialyzed o/n against dialysis buffer. The following day, the dialyzed protein was centrifuged for 15 min at 4,000 × *g* and 4 °C to remove any aggregates, and then concentrated using an Amicon® 100 kDa MWCO Ultra Centrifugal Filter (cellulose pore size 100 kDa nominal molecular weight limit; Millipore) for Rubisco proteins. For 6PGD, an Amicon® 10 kDa MWCO Ultra Centrifugal Filter (Millipore) was used. Protein concentrations were calculated based on *A*_280_ measured on a NanoDrop using extinction coefficients calculated based on their individual amino acid contents to obtain the molar concentration (**Supplementary Table 2**). 100 µL aliquots of concentrated protein were flash-frozen in liquid N_2_ and stored at −80 °C.

### Purification of Rubisco from leaf extracts

Healthy, fully expanded leaves from *Arabidopsis thaliana* and *Spinacia oleracea* were harvested and frozen in liquid N_2_. Frozen leaves were homogenized in 10 mL of extraction buffer (100 mM EPPS at pH 8.1, 15 mM MgCl_2_, 1 mM EDTA, 5 mM DTT, 1% PVPP) using a 40 mL glass homogenizer (Wheaton) on ice. The total homogenate was centrifuged at 20,000 × *g* at 4 °C for 20 min. The supernatant was collected and filtered using a 0.22 µM filter to remove any remaining leaf particulate matter. Clarified sample was loaded onto a MonoQ 10/100 GL Ion Exchange (IEX) Column (Cytiva) equilibrated with IEX buffer A (50 mM Tris-HCl at pH 8.0, 10 mM NaCl). The protein was eluted with a linear salt gradient (0.01–0.5 M NaCl) over 10 column volumes using IEX buffer B (50 mM Tris-HCl at pH 8.0, 1 M NaCl). Protein eluting at ∼0.3 M NaCl was collected and concentrated using an Amicon® 100 kDa filter at 4,000 × *g* at 4 °C to ∼0.5 mL before being loaded on an Enrich SEC 650 gel filtration column (Biorad) equilibrated with SEC buffer (100 mM EPPS at pH 8.1, 15 mM MgCl_2_, 1 mM EDTA) and eluted at 0.5 mL/min. Fractions containing Rubisco were identified by their abundant *A*_280_ profile and pooled, then further concentrated with an Amicon® 100 kDa filter at 4 °C to a final concentration of ∼3 mg/mL. Protein concentrations were calculated based on *A*_280_ measured on a NanoDrop using extinction coefficients, as previously described (**SI Table 2**). Samples were aliquoted and flash-frozen in liquid N_2_ and stored at −80 °C.

### RubiSpec-MS assay protocol

#### Sample preparation

Reactions were set up as described previously (Kane *et al*., 1994) for the Radiolabeled RuBP Method, but the protocol is also provided in detail here for clarity (**Fig. 2**). We note that, for validation, at least one sample in every set of reactions should contain a reliable Rubisco of known *S*_C/O_ appropriate for the gas mixture being used. Reactions were performed in septum-sealed 20 mL glass vials (ChemGlass Life Sciences) containing 20 µL of 10 mg/mL carbonic anhydrase, 10–20 µL of purified Rubisco (40 µM), and *S*_C/O_ assay buffer (30 mM triethanolamine adjusted to pH 8.30 with acetic acid, 15 mM Mg(OAc)_2_) at a final volume of 980 µL. The reactions were equilibrated as open systems under a defined atmosphere of O_2_ and CO_2_. The open system was maintained via two 21-gauge syringe needles inserted into the septa of each vial. To ensure sufficient positive pressure while also avoiding evaporation of the reaction mixture over the course of the assay, the inlet needle was briefly immersed in Milli-Q H_2_O and the gas flow rate was adjusted to maintain a steady stream of bubbles. The vent needle was inserted through the vial septa, followed by the gas needle. The needles were positioned such that they did not touch the reaction mixture in the vial but were also deep enough to prevent the needles detaching from the septa during water bath shaking. Based on the anticipated range for the *S*_C/O_ value, different gas mixtures of O_2_ and CO_2_ were employed. For plant Rubisco enzymes (with a predicted *S*_C/O_ of 60–100), a gas mixture of 999,500 ppm O_2_ and 500 ppm CO_2_ (Linde) was used. For bacterial Rubisco enzymes (with a predicted specificity of 10–60), a gas mixture of 998,000 ppm O_2_ and 2000 ppm CO_2_ (Linde) was used. Reaction vials were placed in a water bath held at 25.0 °C, as measured by an infrared thermometer, and left to shake at 100 RPM for at least 30 min for gas and temperature equilibration. 20 µL of 30 mM RuBP was injected using a gas-tight LC-MS syringe (Hamilton), bringing the final reaction mixture volume to 1 mL. Reactions were then allowed to proceed for 1 h in the shaking water bath at 100 RPM at 25.0 °C. Immediately after the 1 h reaction period, each sample was passed through an Amicon® 3□kDa MWCO Ultra Centrifugal Filter (cellulose pore size 3-kDa nominal molecular weight limit; Millipore). 40 µL aliquots of each filtered sample were diluted with 10 µL of 150 mM aqueous NH_4_HCO_3_ at pH 9.00 and 50 µL ACN to yield a final solution of 15 mM NH_4_HCO_3_ at pH 9.00 in H_2_O/ACN 50:50 (v/v). These solutions were transferred to LC vials fitted with 300 µL glass inserts and septum caps (Waters). The remaining filtered sample was flash-frozen in liquid N_2_ and stored at −80 °C for later analysis.

**Figure 2.**
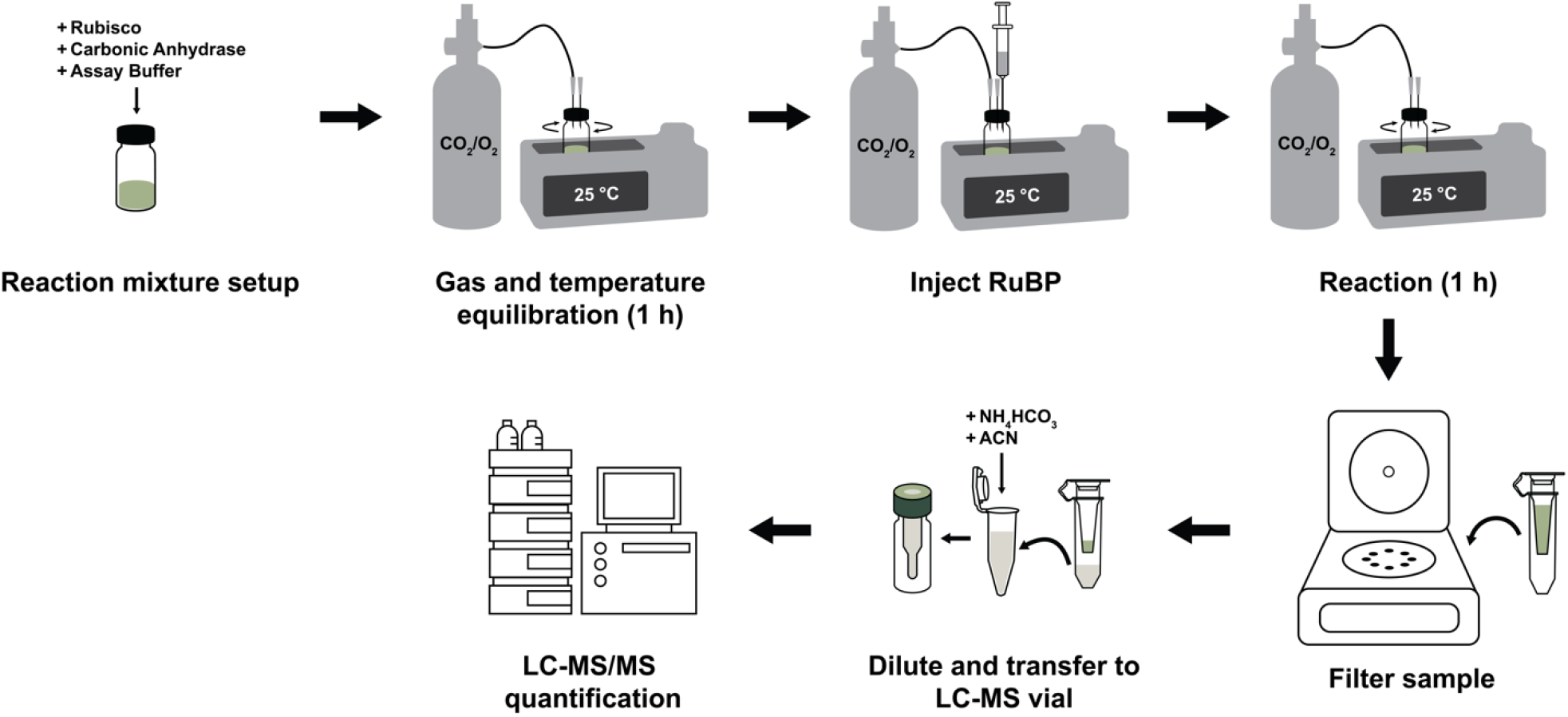
Schematic of the RubiSpec-MS (Rubisco Specificity by LC-MS/MS) method workflow. The reaction set-up and assay framework were adapted from the Radiolabeled RuBP Method (Kane et al., 1994), with key modifications to the reaction clean-up protocol for separation and quantification of non-radiolabeled reaction products using LC-MS/MS. Gas delivery can be split to multiple sample vials using a stainless steel aquarium air flow manifold equipped with adjustable valves. Abbreviation: RuBP (ribulose-1,5-bisphosphate).

#### Standard Preparation

Standards were prepared using commercially available 3PGA, 2PG, and in-house synthesized RuBP. A standard stock containing 1.25 mM each of 3PGA, 2PG, and RuBP was prepared in *S*_C/O_ assay buffer. The stock solution was then serially diluted 1:1 in *S*_C/O_ assay buffer five times to generate a six-point calibration curve. 40 µL aliquots of each standard were diluted with 10 µL of 150 mM NH_4_HCO_3_ at pH 9.00 and 50 µL ACN to final standard concentrations of 500.0, 250.0, 125.0, 62.50, 31.25, and 15.63 µM in 15 mM NH_4_HCO_3_ (pH 9.00) in H_2_O/CAN 50:50 (v/v) The final standard solutions were transferred to LCMS vials fitted with 300 µL glass insert and septum caps (Waters).

#### Metabolite quantification with LC-MS/MS

2PG, 3PGA, and RuBP quantification by LC-MS/MS was performed using a Waters ACQUITY Premier UPLC coupled to a Waters Xevo TQ-S Micro Triple Quadrupole Mass Spectrometer. Samples were run on an Atlantis Premier BEH Z-HILIC column (2.1 × 150 mm, 1.7 µm). The column temperature was maintained at 30 °C and samples were held at 8 °C. Mobile phase A was 15 mM aqueous NH_4_HCO_3_ at pH 9.00. Mobile phase B was a mixture of H_2_O and ACN (10:90, v/v) containing a final concentration of 15 mM NH_4_HCO_3_ at pH 9.00. The following LC gradient was used: A linear ramp of 10 % A to 35 % A for 5 min, a hold at 35 % A for 1 min, a linear ramp down to 10 % A for 0.5 min, and a final hold at 10 % A for 8.5 min for equilibration. The injection volume was 2 µL, the flow rate was 0.5 mL/min, and the total run time was 15 min. The mass spectrometer was operated in negative ionization mode using multiple reaction monitoring (MRM), with a capillary voltage of 1 kV, desolvation temperature of 50 °C, desolvation gas flow of 1000 L/h, cone gas flow of 50 L/h, and source temperature of 150 °C. Diagnostic transitions, collision energies, cone voltages, and dwell times were optimized by direct infusion of standards using the Waters IntelliStart software program (**Table 1**).

**Table 1.**
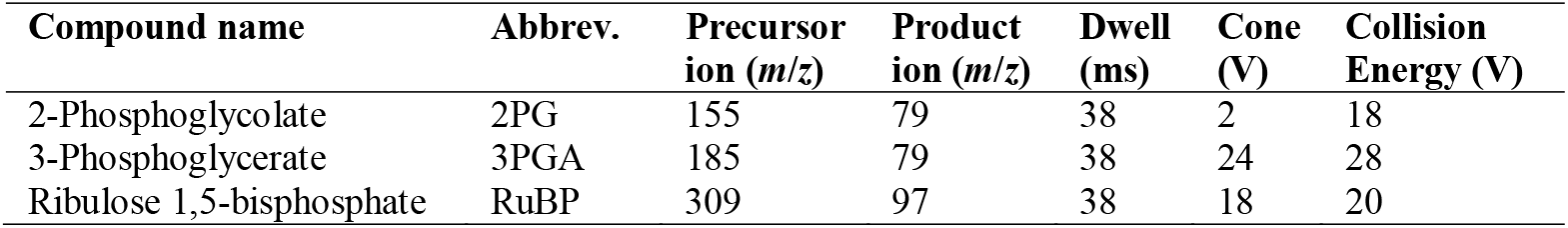
Optimized MRM transition parameters for the quantification of RuBP, 3PGA, and 2PG. Data were acquired in negative electrospray ionization mode, with dwell time, cone voltage, and collision energies tuned to maximize sensitivity for each metabolite.

| Compound name | Abbrev. | Precursor ion (*m/z*) | Product ion (*m/z*) | Dwell (ms) | Cone (V) | Collision Energy (V) |
| --- | --- | --- | --- | --- | --- | --- |
| 2-Phosphoglycolate | 2PG | 155 | 79 | 38 | 2 | 18 |
| 3-Phosphoglycerate | 3PGA | 185 | 79 | 38 | 24 | 28 |
| Ribulose 1,5-bisphosphate | RuBP | 309 | 97 | 38 | 18 | 20 |

**Table 2.** Temperature-dependent gas solubilities of O_2_ and CO_2_ at 1 atm (Bunsen coefficient, unitless) used to convert atmospheric partial pressures to the dissolved concentrations of each gas for determining Rubisco *S*_C/O_. Adapted from (Dawson, 1986).

| Temp (°C) | O<sub>2</sub> | CO<sub>2</sub> | O<sub>2</sub>/CO<sub>2</sub> |
| --- | --- | --- | --- |
| 0 | 0.049 | 1.713 | 0.029 |
| 5 | 0.043 | 1.424 | 0.030 |
| 10 | 0.038 | 1.194 | 0.032 |
| 15 | 0.034 | 1.019 | 0.033 |
| 20 | 0.031 | 0.878 | 0.035 |
| 25 | 0.028 | 0.759 | 0.037 |
| 30 | 0.026 | 0.665 | 0.039 |
| 35 | 0.024 | 0.592 | 0.041 |
| 37 | 0.024 | 0.567 | 0.042 |
| 48 | 0.023 | 0.550 | 0.042 |
| 40 | 0.023 | 0.530 | 0.043 |
| 45 | 0.022 | 0.479 | 0.046 |
| 50 | 0.021 | 0.436 | 0.048 |

Samples and standards were run in triplicate. Peak integration and quantification were performed in Waters MassLynx software, and metabolite concentrations were exported to Microsoft Excel for *S*_C/O_ calculations.

### Specificity Factor (s_C/O_) Calculation

Rubisco *S*_C/O_ is defined as the ratio between the carboxylation efficiency (*V*_C_*/K*_C_) and oxygenation efficiency (*V*_O_*/K*_O_), where *V*_C_ and *V*_O_ represent the maximum velocities of carboxylation and oxygenation, respectively, and *K*_C_ and *K*_O_ represent the Michaelis–Menten constants for CO_2_ and O_2_, respectively (Laing *et al*., 1974).

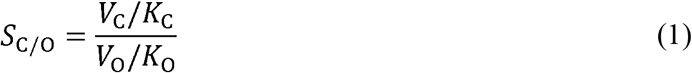

O_2_ and CO_2_ are competitive inhibitors of carboxylation and oxygenation, respectively, thus **Equation 1** can be rewritten as:

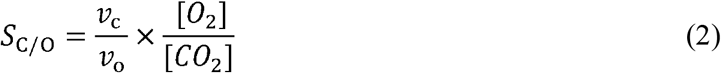

where *v_c_* represents velocity of carboxylation, *v_O_* represents the velocity of oxygenation, and [*O*_2_] and [*CO*_2_] represent the concentration of O_2_ and CO_2_, respectively (Laing *et al*., 1974). From **Equation 2**, *S*_C/O_ can also be defined as the number of carboxylation events per oxygenation event at equal partial pressures of dissolved CO_2_ and O_2_ (Kane *et al*., 1994; Kostov & McFadden, 1995; Von Caemmerer, 2000). The ratio of the carboxylation to oxygenation can then be approximated by the following equation:

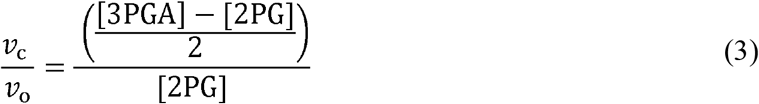

where [3PGA] and [2PG] represent the concentrations of 3PGA and 2PG, respectively. The relative concentrations of dissolved O_2_ and CO_2_ are calculated by multiplying their partial pressures (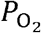 and 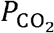) in the gas mixture delivered to the reaction vial by their respective temperature-dependent solubility constants at a specified temperature and pressure (represented by *α*_O_ and *α*_C_ for O_2_ and CO_2_, respectively).

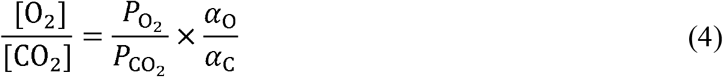

Substituting **Equations 3** and **4** into **Equation 2** yields **Equation 5**:

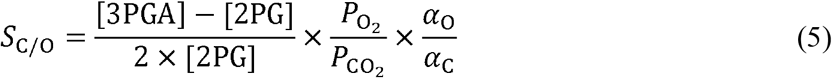

The solubility constants at varying temperatures are provided in **Table 2** or can be calculated from Dawson (Dawson, 1986). Given the known molar gas ratio of O_2_ and CO_2_, the RubiSpec-MS method can be used to determine *S*_C/O_ across a range of temperatures from **Equation 5**. At 25 °C and 1 atm, the temperature-dependent solubility constants of O_2_ and CO_2_ are 0.028 and 0.759, respectively (Dawson, 1986). For the RubiSpec-MS method, *S*_C/O_ at 25 °C is then calculated from metabolite quantification by LC-MS/MS using **Equation 6**:

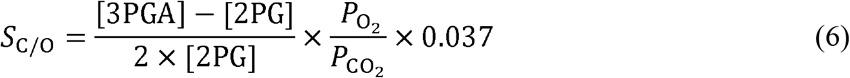

where the constant 0.037 is the ratio of the temperature-dependent solubility constants of O_2_ and CO_2_ at 25 °C (**Table 2**). The *S*_C/O_ of each reaction sample was averaged across triplicate LC-MS/MS runs. Then, the calculated *S*_C/O_ value for each Rubisco was averaged across biological replicates and reported as mean ± S.D.

### Method Validation

A simulated sample matrix was prepared and filtered with the same method described above for *S*_C/O_ measurement, except for the injection of RuBP. Then, a standard mixture of RuBP, 3PGA, and 2PG was spiked into the matrix to achieve a final analyte concentration of 250 μM. Matrix effect was calculated by **Equation 7**:

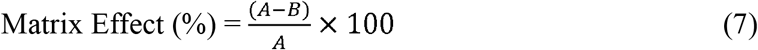

where A is the peak area of the analyte in neat solvents and B is the peak area of the analyte in a sample prepared according to the RubiSpec-MS assay protocol with no RuBP injection, then later spiked with analytes to the same standard concentration (Zhou *et al*., 2017). A 250 μM standard mix was injected 15 times on the same day to assess intraday precision of the method. The limit of detection (LOD) and limit of quantification (LOQ), reported by MassLynx software, were calculated by multiplying the ratio of the standard deviation of the response to the slope of the calibration curve by 3 and 10, respectively.

## RESULTS

### LC-MS/MS method development and optimization

To develop a faster, safer, and more efficient method to measure Rubisco *S*_C/O_ without relying on radiolabeled materials, we first focused on the separation and quantification of 2PG, 3PGA, and RuBP with LC-MS/MS. We were motivated by prior work showing the successful separation of sugar phosphates using hydrophilic interaction liquid chromatography (HILIC) columns under high-pH mobile phase conditions, followed by detection and quantification by tandem mass spectrometry (MS/MS) (Smith *et al*., 2021; Su *et al*., 2023; Serafimov & Lämmerhofer, 2024). We began by evaluating the detection of commercially available 2PG, 3PGA, and RuBP via direct infusion of a combined standard solution containing all three compounds on the Waters Xevo TQ-S Micro Triple Quadrupole Mass Spectrometer. We optimized multiple reaction monitoring (MRM) transitions and MS/MS parameters using the IntelliStart software, then selected deprotonated molecular ions [M-H]^-^ as the precursor ions for all compounds (**Table 1**). We selected the most abundant fragments for quantification as the product ions, and the second-most abundant fragments for qualification.

Following successful detection of these compounds, we tested chromatographic separation on the Atlantis Premier BEH Z-HILIC Column. We prepared mobile phases fresh for every batch run, with a pH accuracy of ± 0.01 units to ensure reproducibility (Smith *et al*., 2021). 2PG, 3PGA, and RuBP eluted at 4.05, 4.32, and 4.80 min, respectively, demonstrating effective separation (**Table 3** and **Fig. 3**). Linear regression over a concentration range of 15.63–500.0 µM yielded correlation coefficients of around 0.97, indicating reliable quantification (**Fig. 3**). The LOD and LOQ, calculated using the MassLynx software, are reported in **Table 3**. The intraday precision for each analyte was <15%, meeting accepted analytical criteria (Whitmire *et al*., 2011). We observed a positive matrix effect on 2PG and RuBP, indicating that the sample matrix suppressed ion detection of both analytes. We observed a negative matrix effect on 3PGA, indicating that the sample matrix enhanced its ion detection. All matrix effect values were within the acceptable ± 20% range (**Table 3**) (Zhou *et al*., 2017).

**Table 3.** Method validation results. Matrix effect was calculated by comparing the peak area of an analyte in neat solvents and the peak area of the same analyte spiked into a simulated matrix at the same concentration (250 µM), *n = 5*. Intraday precision was calculated from injecting the same standard mix (250 µM) 15 × on the same day. Relative standard deviation (RSD) % = (SD × 100)/mean. The limit of detection (LOD) and limit of quantification (LOQ), calculated using the MassLynx software, were obtained by multiplying the ratio of the standard deviation of the response to the slope of the calibration curve by 3 and 10, respectively.

| Compound name | Abbrev. | Retention time (min) | Matrix effect (%) | Intraday precision (RSD%) | LOD ($\mu\text{M}$) | LOQ ($\mu\text{M}$) |
| --- | --- | --- | --- | --- | --- | --- |
| 2-Phosphoglycolate | 2PG | 4.05 | 9.12 | 12.65 | 9.98 | 10.02 |
| 3-Phosphoglycerate | 3PGA | 4.32 | -6.80 | 5.69 | 1.98 | 4.08 |
| Ribulose-1,5-bisphosphate | RuBP | 4.80 | 9.36 | 7.78 | 3.49 | 4.54 |

**Figure 3.**
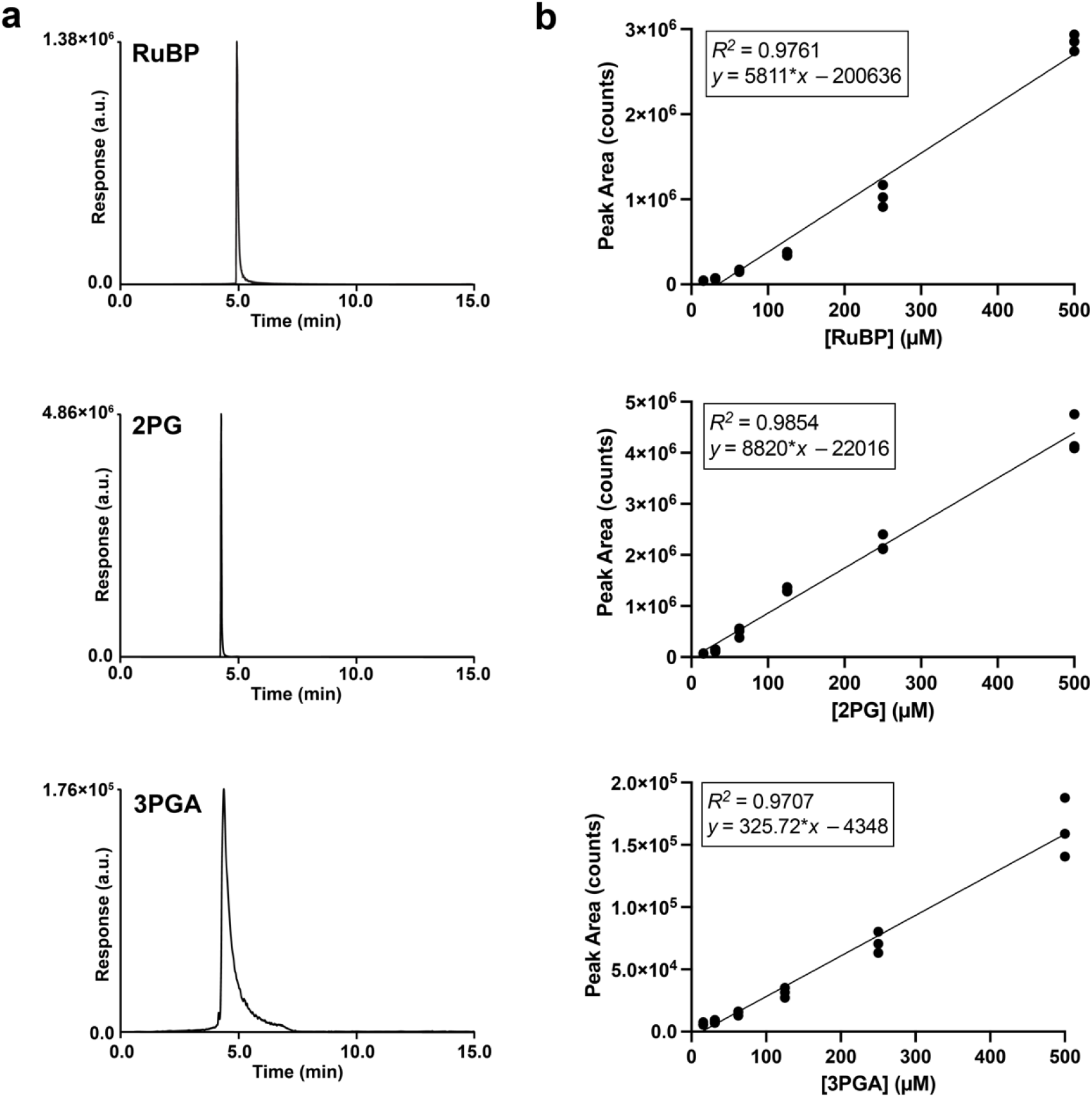
LC-MS/MS characterization of RuBP, 2PG, and 3PGA. **(a)** Representative extracted ion chromatograms demonstrating peak resolution and retention times for RuBP, 2PG, and 3PGA. **(b)** Linear calibration curves of RuBP, 2PG, and 3PGA across a 15.63–500.0 µM range. Abbreviations: RuBP (ribulose-1,5-bisphosphate), 2PG (2-phosphoglycolate), 3PGA (3-phosphoglycerate).

### Adaptation of *S*_C/O_ assay from established radiometric methods

Upon successful quantification of commercial RuBP, 2PG, and 3PGA, we aimed to quantify the total 2PG and 3PGA produced by Rubisco catalysis and determine if the known *S*_C/O_ for various model Rubisco enzymes could be arrived at accurately using this approach and **Equation 6**. For coherency with prior measurements, the technical aspects of sample preparation and clean-up in the RubiSpec-MS method were adapted from the Radiolabeled RuBP Method (Kane *et al*., 1994), with minor modifications to the reaction set-up and assay workflow to increase efficiency. We retained similar buffer conditions and reaction mixtures. Critically, we also maintained the inclusion of carbonic anhydrase to ensure rapid interconversion of CO_2_ and HCO_3_. We also maintained a pH of 8.30 (within the range of 7.4–8.9, where previous studies have shown no significant impact of pH on Rubisco *S*_C/O_) (Jordan & Ogren, 1984; Badger & Price, 1994). Given optimized metabolite quantification over the 15.63–500.0 µM range and requisite dilution postreaction with ACN and NH_4_HCO_3_ for LC-MS/MS analysis, the RubiSpec-MS assay used 20 µL of 30 mM RuBP (final concentration of 600 µM in the 1-mL reaction mixture) to generate sufficiently strong signal for detection of the resulting 3PGA and 2PG. 600 µM RuBP is well above the Michaelis-Menten constant for RuBP (*K*_RuBP_), which ranges from 0.1–100 µM (Flamholz *et al*., 2019). Previous studies determined *S*_C/O_ is independent of RuBP concentration at saturating RuBP concentrations relative to *K*_RuBP_, because the catalytic rates are not limited by substrate availability (Jordan & Ogren, 1984). Only in the case where RuBP concentration is low enough that *k*, approaches zero would *K*_RuBP_ become a critical parameter. Thus, varying concentrations of RuBP ≥ 100 µM can be used depending on the LOQ and quantification range of the MS employed.

Consistent gas ratios of CO_2_ and O_2_ must be maintained over the course of the Rubisco reaction, as the dissolved CO_2_ and O_2_ concentrations directly relate to the derivatization of *S*_C/O_. The Radiolabeled RuBP Method employs an open gas system, where a fixed ratio of CO_2_ and O_2_ are delivered to a septa-sealed reaction vial fitted with a vent needle, allowing for constant gas equilibration. Historically, the need for precision control over gas mixtures used in such an open system required the use of Wösthoff gas mixing pumps (Wösthoff Messtechnik GmbH; Bochum, Germany) for accurate mixing of pure gases (Mueller-Cajar *et al*., 2007; Kubien *et al*., 2008). However, precision gas mixtures can now be commonly purchased from commercial gas suppliers and are routinely used for *S*_C/O_ measurements via the Radiolabeled RuBP Method (Davidi *et al*., 2020; Schulz *et al*., 2022). For RubiSpec-MS, we also used these gas mixtures, ranging from 0.05–0.2 % CO_2_ depending on the expected *S*_C/O_ of the Rubisco enzyme under study. Due to the higher specificity factors of plant Rubiscos, a lower CO_2_ concentration was necessary to generate sufficient 2PG in the reaction product mixture for accurate quantification. We used a gas mixture of 0.05% CO_2_ and 99.95% O_2_ to measure *S*_C/O_ of plant Rubiscos and a gas mixture of 0.2% CO_2_ and 99.8% O_2_ to measure *S*_C/O_ of bacterial Rubiscos.

Maintenance of stable temperature over the duration of the Rubisco reaction is of critical importance, as the temperature dependence of *S*_C/O_ and CO_2_/O_2_ solubility are well-documented (Jordan & Ogren, 1984; Galmés *et al*., 2005; Yamori *et al*., 2006; Hermida-Carrera *et al*., 2016). We performed all *S*_C/O_ measurements at 25.0 °C, the standard temperature for Rubisco kinetic measurements, to enable robust comparison with literature values. To ensure this temperature, we regularly checked the water bath every ∼15 min with an infrared temperature gun thermometer, throughout Rubisco activation and reaction.

To ensure that the generated reaction products fell within the linear range of the RubiSpec-MS calibration curve, the initial substrate concentrations and subsequent sample dilution steps were considered. Both carboxylation and oxygenation of RuBP yields two product molecules, thus a RuBP concentration of 600 μM in the 1-mL reaction mixture could yield a maximum of 600 μM of 2PG if only oxygenation reactions occurred, or 1.2 mM of 3PGA if only carboxylation reactions occurred. After a 0.4× dilution of the reaction mixture prior to LC-MS/MS analysis, the maximum theoretical product concentration is reduced to 240 μM and 480 μM for 2PG and 3PGA, respectively, falling within the demonstrated analytical range.

The main steps where RubiSpec-MS deviates significantly from the Radiolabeled RuBP Method lie in the product clean-up and quantification. In the Radiolabeled RuBP Method, alkaline phosphatase is added following the reaction to cleave the phosphate groups from the unreacted RuBP and reaction products, leaving radiolabeled ribulose, phosphoglycerate, and phosphoglycolate. Dephosphorylated products are then applied to an anion-exchange resin, which binds to the phosphate groups and allows ribulose to pass through. Removal of ribulose is necessary in the Radiolabeled RuBP Method because ribulose coelutes with glycerate during high-performance liquid chromatography (HPLC). The radiolabeled glycerate and glycolate are then eluted from the resin with acid and subsequent HPLC separates the labeled compounds in the reaction mixture. Finally, fractions of the elution are collected for liquid scintillation count-ing and *S*_C/O_ is calculated from the ratio of radiolabeled glycerate and glycolate formed under a given atmosphere (**Fig. 1c**) (Kane *et al*., 1994).

In contrast to the Radiolabeled RuBP Method, successful separation of the direct reaction products using LC-MS/MS eliminates the need for dephosphorylation by alkaline phosphatase and removal of ribulose via anion-exchange resin prior to liquid chromatography. Instead, we allowed the Rubisco reaction to proceed for 1 h after RuBP injection to ensure the reaction reached completion, then immediately passed the sample through a 3□kDa MWCO filter to remove proteins and minimize contamination of the HILIC column. Finally, we diluted the filtered sample to a final concentration of 15 mM NH_4_HCO_3_ (pH 9.00) in H_2_O/ACN 50:50 (v/v) for LC-MS/MS analysis (**Fig. 2**). *S*_C/O_ was then calculated from the quantification of unlabeled 3PGA and 2PG formed under a given atmosphere according to the equation in **Fig. 1e**.

### RubiSpec-MS Method Validation and Application

After method optimization and modification of the Radiolabeled RuBP Method workflow to suit LC-MS/MS analysis, we next applied the RubiSpec-MS method to measure the *S*_C/O_ of Rubisco enzymes commonly used as references or that have otherwise had their *S*_C/O_ parameter reported in the literature. We measured the *S*_C/O_ of these representative bacterial and plant Rubiscos using RubiSpec-MS in parallel with the previously described Radiolabeled RuBP Method, with the modification that [1-^3^H]-RuBP was used instead of [1-^14^C]-RuBP (Kane *et al*., 1994). In each sample batch prepared, we included a Rubisco enzyme with a well-documented *S*_C/O_ value as an internal positive control to ensure consistent system conditions (*Nt*LS for measurement of higher plant Rubiscos with higher *S*_C/O_ values and RbcM for measurement of Rubisco with lower *S*_C/O_ values). All Rubiscos returned *S*_C/O_ values consistent with parallel samples run with the Radiolabeled RuBP method within a 95% confidence interval. Moreover, *S*_C/O_ values measured with both the RubiSpec-MS method and the Radiolabeled RuBP method aligned with those previously reported in the literature (**Table 4**).

**Table 4.** *S*_C/O_ values determined using the LC-MS/MS method closely correlated with other routinely used methods. Values for this study are reported as mean ± S.D. (*n* = 3–6). Specificity factors determined using the radiolabeled RuBP method were sourced from the literature for *R. rubrum* (Kane *et al*., 1994), *S. elongatus* PCC 6301 (Kane *et al*., 1994), *C. sphaeroides* (Zhou *et al*., 2023), *A. thaliana* (Whitney *et al*., 2015), *N. tabacum* (Kane *et al*., 1994), and *S. oleracea* (Kane *et al*., 1994). *S*_C/O_ values determined using the O_2_ electrode method were sourced from literature for *R. rubrum* (Parry *et al*., 1989), *S. elongatus* PCC 6301 (Parry *et al*., 1989), *N. tabacum* (Parry *et al*., 1989), and *S. oleracea* (Hermida-Carrera *et al*., 2016).

| Species | Organism | Form | RubiSpec-MS | Radiolabeled RuBP (this work) | Radiolabeled RuBP (lit.) | $O_2$ electrode (lit.) |
| --- | --- | --- | --- | --- | --- | --- |
| *Rhodospirillum rubrum* | Proteobacteria | II | 11.9 ± 1.0 | 11.5 ± 0.3 | 12.3 ± 0.1 | 9 ± 0.9 |
| *Synechococcus elongatus* PCC 6301 | Cyanobacteria | IB | 38.8 ± 3.4 | 39.0 ± 2.4 | 42.5 ± 1.0 | 52 ± 4.2 |
| *Cereibacter sphaeroides* | Purple bacteria | IC | 63.4 ± 4.1 | 62.1 ± 2.2 | 59.5 ± 0.6 | - |
| *Arabidopsis thaliana* | Plant | IB | 84.7 ± 2.4 | 86.2 ± 0.9 | 80 ± 2 | - |
| *Nicotiana tabacum* | Plant | IB | 80.3 ± 3.3 | 82.0 ± 1.8 | 81.9 ± 1.0 | 89 ± 4.1 |
| *Spinacea oleracea* | Plant | IB | 83.9 ± 3.7 | 90.6 ± 4.1 | 82.1 ± 0.8 | 97.0 ± 1.2* |

\*Reported as SE instead of SD.

### Synthesized RuBP provides more accurate specificity factor measurements

While a primary advantage of the RubiSpec-MS method is the elimination of radiolabeled RuBP, accuracy of *S*_C/O_ determination still relies on substrate purity. Previous studies found that commercial non-pure RuBP yields poor kinetic measurements due to inhibitory impurities (Sharwood *et al*., 2016). To evaluate whether the RubiSpec-MS method is similarly sensitive to these impurities, we benchmarked the performance of commercial RuBP against in-house synthesized RuBP. For RbcM and *Syn*LS, the *S*_C/O_ values obtained using commercial RuBP were comparable to those derived from synthesized RuBP within a 95% confidence interval. However, the *S*_C/O_ measurements for *Cs*LS exhibited a higher standard deviation when using the commercial substrate (**Table 5**). Thus, while the RubiSpec-MS method is compatible with commercial RuBP, the use of in-house synthesized RuBP is still recommended to achieve maximum precision and consistency in *S*_C/O_ determinations.

**Table 5.** *S*_C/O_ determination using commercial and synthesized RuBP. Commercial RuBP yields *S*_C/O_ values comparable to synthesized RuBP within a 95% confidence interval, though in-house synthesized substrate remains optimal for maximizing reproducibility and minimizing variant-specific standard deviations. Values are reported as mean ± S.D (*n* = 3–6).

| Species | Commercial RuBP | Synthesized RuBP |
| --- | --- | --- |
| *Rhodospirillum rubrum* | 11.9 ± 1.0 | 11.8 ± 0.3 |
| *Synechococcus elongatus* PCC 6301 | 41.0 ± 2.9 | 38.8 ± 3.4 |
| *Cereibacter sphaeroides* | 61.3 ± 8.9 | 63.4 ± 4.1 |

## DISCUSSION

Accurate quantification of Rubisco *S*_C/O_ is essential to studying the catalytic mechanism of Rubisco and engineering improved Rubisco enzymes. Current benchmark methods for *S*_C/O_ quantification rely on synthesis of radioactive materials that cannot be commercially obtained, and necessitate bespoke equipment dedicated for radioactive materials separation, measurement, handling, and storage. Moreover, these methods require personnel trained for safe handling and disposal of radioactive materials. The complexity and time-intensive demands of current methodologies have limited large-scale measurement of *S*_C/O_ in natural and mutant Rubiscos, resulting in significant knowledge gaps. Here, we present a faster, more accessible method to quantify *S*_C/O_ without radiolabeled materials and review its performance relative to current standard methods.

The most widespread method of Rubisco *S*_C/O_ quantification, the O_2_ Electrode Method, requires a closed system to accurately quantify O_2_ consumption by Rubisco. However, the decrease in O_2_ concentration during the reaction may limit oxygenase activity, which is disadvantageous for measuring the slow oxygenation rates in higher plant Rubisco enzymes. The dissolved CO_2_ concentration is maintained by the presence of carbonic anhydrase in the buffer, and thus the change in CO_2_/O_2_ ratio results in a significantly higher *S*_C/O_ measurement in plant Rubisco enzymes by the O_2_ Electrode Method in comparison to the Radiolabeled RuBP Method (Iñiguez *et al*., 2021). While the Radiolabeled RuBP Method provides accurate control of the CO_2_/O_2_ gas ratio, it demands a more complex and time-intensive protocol to accurately separate and quantify the radiolabeled products.

To overcome these limitations of existing Rubisco *S*_C/O_ assays, we developed a simpler, more efficient assay free of radioactive materials. The RubiSpec-MS method for determining *S*_C/O_ adopts the same open gas system and reaction set-up of the Radiolabeled RuBP Method but leverages LC-MS/MS to separate and quantify the direct reaction products, eliminating the needs for dephosphorylation by alkaline phosphatase and removal of ribulose via anion-exchange resin prior to liquid chromatography. The omission of reaction cleanup steps and scintillation counting of microtiter plates, in addition to a shorter LC gradient (15 min for the RubiSpec-MS method vs. 22 min for the Radiolabeled RuBP Method), reduces the analysis time per sample and simplifies the assay workflow. From reaction set-up to *S*_C/O_ calculation, the RubiSpec-MS protocol can be completed in roughly half the time of the Radiolabeled RuBP Method. Moreover, many samples can be prepared in batches and quantified on the LC-MS/MS without researcher monitoring, allowing numerous Rubiscos to be assayed in parallel.

While radiolabeled assays offer high analytical sensitivity, their complexity and limited throughput have prevented large-scale analysis of Rubisco diversity, unlike measurements of other Rubisco kinetic parameters, such as carboxylation rate (Davidi *et al*., 2020). RubiSpec-MS enables parallelized measurements across multiple Rubisco variants and species. Although it is still recommended to synthesize unlabeled RuBP to avoid \ contaminants present in current commercial preparations and maximize reproducible *S*_C/O_ measurements, this requirement is substantially less restrictive than radioactive substrate synthesis and handling.

In sum, RubiSpec-MS avoids restrictions of radioactive assays and is suitable for use on shared mass spectrometry instruments, making *S*_C/O_ measurement feasible for research groups that rely on LC-MS machines in departmental core facilities. The RubiSpec-MS assay produced *S*_C/O_ values consistent with values measured with the established Radiolabeled RuBP Method as well as available in prior literature, demonstrating reproducible, accurate measurement of *S*_C/O_. Given its ease of implementation, parallelizability, and equivalent accuracy to previous radiometric methods, RubiSpec-MS is expected to make an accessible and accurate *S*_C/O_ assay available to a broad range of Rubisco researchers.

## ACKNOWLEDGEMENTS

We thank Professor Mary Gehring (MIT) for providing *A. thaliana* leaf tissue and Professor Spencer Whitney (ANU) for providing the plasmids pTrc-*Syn*LS, pCDF-NtAsmbl, pET16-*Nt*LS*Rca*, and pUSP2-^His6^USP2. We also thank Dr. Terri Sosienski, Dr. Alexandre F. Gomes, and Paul Kowalski of Waters^™^ for helpful advice on the use of the Atlantis Premier BEH Z-HILIC column and the Waters Xevo TQ-S Micro Triple Quadrupole Mass Spectrometer.

## AUTHOR CONTRIBUTIONS

R.H.W. conceptualized the use of LC-MS/MS to determine Rubisco *S*_C/O_. A.W., R.H.W., and Y.Z. developed the LC-MS/MS method and performed experiments. M. K. provided helpful advice on LC-MS/MS analysis. A.W., R.H.W., Y.Z., and J.L.M. performed the synthesis and purification of reagents and proteins. All authors designed experiments and analyzed data. A.W., R.H.W., and M.D.S. drafted the manuscript. All authors edited the manuscript.

## DATA AVAILABILITY

The plasmid sequences and purified protein concentrations used in this study are provided in **Supplementary Tables 1** and **2**, respectively. The raw data for the matrix effect and intraday precision calculations are provided in **Supplementary Tables 3** and **4**, respectively. The data for *S*_C/O_ determination by the RubiSpec-MS method are provided in **Supplementary Table 5**. The data for *S*_C/O_ determination by the Radiolabeled RuBP method are provided in **Supplementary Table 6**.

## REFERENCES

Aguiló-Nicolau P, Iñiguez C, Capó-Bauçà S, Galmés J. 2024. Rubisco kinetic adaptations to extreme environments. Plant J. 119: 2599–2608.

Badger MR, Price GD. 1994. The role of carbonic anhydrase in photosynthesis. Annu. Rev. Plant Biol. 45: 369–392.

Bernacchi C, Singsaas E, Pimentel C, Portis Jr A, Long SP. 2001. Improved temperature response functions for models of RubiscoLlimited photosynthesis. Plant Cell Environ. 24: 253–259.

Bowes G. 1991. Growth at elevated CO2: photosynthetic responses mediated through Rubisco. Plant Cell Environ. 14: 795–806.

Brooks A, Farquhar G. 1985. Effect of temperature on the CO2/O2 specificity of ribulose-1,5-bisphosphate carboxylase/oxygenase and the rate of respiration in the light: estimates from gas-exchange measurements on spinach. Planta 165: 397–406.

Buck S, Rhodes T, Gionfriddo M, Skinner T, Yuan D, Birch R, Kapralov MV, Whitney SM. 2022. Escherichia coli expressing chloroplast chaperones as a proxy to test heterologous Rubisco production in leaves. J. Exp. Bot. 74: 664–676.

Busch FA. 2020. Photorespiration in the context of Rubisco biochemistry, CO2 diffusion and metabolism. Plant J. 101: 919–939.

Catanzariti A-M, Soboleva TA, Jans DA, Board PG, Baker RT. 2004. An efficient system for high-level expression and easy purification of authentic recombinant proteins. Protein Sci. 13: 1331–1339.

Chen T, Riaz S, Davey P, Zhao Z, Sun Y, Dykes GF, Zhou F, Hartwell J, Lawson T, Nixon PJ, et al. 2022. Producing fast and active Rubisco in tobacco to enhance photosynthesis. Plant Cell 35: 795–807.

Cousins AB, Ghannoum O, Von Caemmerer S, Badger M. 2010. Simultaneous determination of Rubisco carboxylase and oxygenase kinetic parameters in Triticum aestivum and Zea mays using membrane inlet mass spectrometry. Plant Cell Environ. 33: 444–452.

Davidi D, Shamshoum M, Guo Z, BarDOn YM, Prywes N, Oz A, Jablonska J, Flamholz A, Wernick DG, Antonovsky N, et al. 2020. Highly active Rubiscos discovered by systematic interrogation of natural sequence diversity. EMBO J. 39: e104081.

Dawson RMC. 1986. Data for Biochemical Research: Clarendon Press.

Farquhar GD, von Caemmerer S, Berry JA. 1980. A biochemical model of photosynthetic CO2 assimilation in leaves of C3 species. Planta 149: 78–90.

Flamholz AI, Prywes N, Moran U, Davidi D, Bar-On YM, Oltrogge LM, Alves R, Savage D, Milo R. 2019. Revisiting trade-offs between Rubisco kinetic parameters. Biochemistry 58: 3365–3376.

Frank J, Müh F. 2025. Calculation of specificity constants Γ for Rubisco of different species from microcalorimetric data. Photosynth. Res. 163: 63.

Galmés J, Aranjuelo I, Medrano H, Flexas J. 2013. Variation in Rubisco content and activity under variable climatic factors. Photosynth. Res. 117: 73–90.

Galmés J, Flexas J, Keys AJ, Cifre J, Mitchell RAC, Madgwick PJ, Haslam RP, Medrano H, Parry MAJ. 2005. Rubisco specificity factor tends to be larger in plant species from drier habitats and in species with persistent leaves. Plant Cell Environ. 28: 571–579.

Gionfriddo M, Birch R, Rhodes T, Buck S, Skinner T, Andersson I, Whitney S. 2025. Laboratory evolution of Rubisco solubility and catalytic switches to enhance plant productivity. Nat. Plants 11: 1939–1950.

Hermida-Carrera C, Kapralov MV, Galmés J. 2016. Rubisco catalytic properties and temperature response in crops. Plant Physiol. 171: 2549–2561.

Iñiguez C, Capó-Bauçà S, Niinemets Ü, Stoll H, Aguiló-Nicolau P, Galmés J. 2020. Evolutionary trends in Rubisco kinetics and their co-evolution with CO2 concentrating mechanisms. Plant J. 101: 897–918.

Iñiguez C, Niinemets Ü, Mark K, Galmés J. 2021. Analyzing the causes of method-to-method variability among Rubisco kinetic traits: from the first to the current measurements. J. Exp. Bot. 72: 7846–7862.

Iqbal WA, Lisitsa A, Kapralov MV. 2023. Predicting plant Rubisco kinetics from RbcL sequence data using machine learning. J. Exp. Bot. 74: 638–650.

Jordan DB, Ogren WL. 1984. The CO2/O2 specificity of ribulose-1,5-bisphosphate carboxylase/oxygenase. Planta 161: 308–313.

Kane HJ, Viil J, Entsch B, Paul K, Morell MK, Andrews TJ. 1994. An improved method for measuring the CO2/O2 specificity of ribulose-bisphosphate carboxylase-oxygenase. Aust. J. Plant Physiol. 21: 449–461.

Kane HJ, Wilkin J-M, Portis AR, Jr., John Andrews T. 1998. Potent inhibition of ribulose-bisphosphate carboxylase by an oxidized impurity in ribulose-1,5-bisphosphate. Plant Physiol. 117: 1059–1069.

Kostov RV, McFadden BA. 1995. A sensitive, simultaneous analysis of ribulose-1,5-bisphosphate carboxylase/oxygenase efficiencies: Graphical determination of the CO2/O2 specificity factor. Photosynth. Res. 43: 57–66.

Kubien DS, Brown CM, Kane HJ 2011. Quantifying the amount and activity of Rubisco in leaves. In: Carpentier R ed. Photosynth. Res. Protoc. Totowa, NJ: Humana Press, 349–362.

Kubien DS, Whitney SM, Moore PV, Jesson LK. 2008. The biochemistry of Rubisco in Flaveria. J. Exp. Bot. 59: 1767–1777.

Laing WA, Ogren WL, Hageman RH. 1974. Regulation of soybean net photosynthetic CO2 fixation by the interaction of CO2, O2, and ribulose-1,5-diphosphate carboxylase. Plant Physiol. 54: 678–685.

Long SP, Zhu X-G, Naidu SL, Ort DR. 2006. Can improvement in photosynthesis increase crop yields? Plant Cell Environ. 29: 315–330.

McDonald JL, Shapiro NP, Mengiste AA, Kaines S, Whitney SM, Wilson RH, Shoulders MD. 2025. In vivo directed evolution of an ultrafast Rubisco from a semianaerobic environment imparts oxygen resistance. Proc. Natl. Acad. Sci. 122: e2505083122.

Mueller-Cajar O, Morell M, Whitney SM. 2007. Directed evolution of Rubisco in Escherichia coli reveals a specificity-determining hydrogen bond in the form II enzyme. Biochemistry 46: 14067–14074.

Mueller-Cajar O, Whitney SM. 2008. Evolving improved Synechococcus Rubisco functional expression in Escherichia coli. Biochem. J. 414: 205–214.

Orr DJ, Alcântara A, Kapralov MV, Andralojc PJ, Carmo-Silva E, Parry MA. 2016. Surveying Rubisco diversity and temperature response to improve crop photosynthetic efficiency. Plant Physiol. 172: 707–717.

Parry M, Keys A, Gutteridge S. 1989. Variation in the specificity factor of C3 higher plant Rubiscos determined by the total consumption of ribulose-P2. J. Exp. Bot. 40: 317–320.

Pierce J, Tolbert NE, Barker R. 1980. Interaction of ribulose-bisphosphate carboxylase/oxygenase with transition-state analogs. Biochemistry 19: 934–942.

Prywes N, Phillips NR, Tuck OT, Valentin-Alvarado LE, Savage DF. 2023. Rubisco function, evolution, and engineering. Annu. Rev. Biochem. 92: 385–410.

Rogers A. 2014. The use and misuse of Vc, max in earth system models. Photosynthesis research 119: 15–29.

Schulz L, Guo Z, Zarzycki J, Steinchen W, Schuller JM, Heimerl T, Prinz S, Mueller-Cajar O, Erb TJ, Hochberg GKA. 2022. Evolution of increased complexity and specificity at the dawn of form I Rubiscos. Science 378: 155–160.

Serafimov K, Lämmerhofer M. 2024. Comprehensive coverage of glycolysis and pentose phosphate metabolic pathways by isomer-selective accurate targeted hydrophilic interaction liiquid chromatography-tandem mass spectrometry assay. Anal. Chem. 96: 17271–17279.

Sharwood RE, Sonawane BV, Ghannoum O, Whitney SM. 2016. Improved analysis of C4 and C3 photosynthesis via refined in vitro assays of their carbon fixation biochemistry. J. Exp. Bot. 67: 3137–3148.

Smith KM, Alkhateeb FL, Brennan K, Rainville PD, Walter TH. 2021. Separation of pentose phosphate pathway, glycolysis, and energy metabolites using an ACQUITY Premier System with an Atlantis Premier BEH Z-HILIC Column. Waters Appl. Note 720007411.

Su M, Serafimov K, Li P, Knappe C, Lämmerhofer M. 2023. Isomer selectivity of one- and two-dimensional approaches of mixed-mode and hydrophilic interaction liquid chromatography coupled to tandem mass spectrometry for sugar phosphates of glycolysis and pentose phosphate pathways. J. Chromatogr. A 1688: 463727.

Valegård K, Andralojc PJ, Haslam RP, Pearce FG, Eriksen GK, Madgwick PJ, Kristoffersen AK, van Lun M, Klein U, Eilertsen HC. 2018. Structural and functional analyses of Rubisco from arctic diatom species reveal unusual posttranslational modifications. J. Biol. Chem. 293: 13033–13043.

Von Caemmerer S. 2000. Biochemical models of leaf photosynthesis: Csiro publishing.

Walker BJ, VanLoocke A, Bernacchi CJ, Ort DR. 2016. The Costs of Photorespiration to Food Production Now and in the Future. Annual Review of Plant Biology 67(Volume 67, 2016): 107–129.

Wang Z-Y, Luo S, Sato K, Kobayashi M, Nozawa T. 1998. Measurements of the CO2/O2 specificity of ribulose-1,5-bisphosphate carboxylase/oxygenase by 31P-and 1H-NMR. Photosynth. Res. 58: 103–109.

Whitmire M, Ammerman J, De Lisio P, Killmer J, Kyle D, Mainstone E, Porter L, Zhang T. 2011. LC-MS/MS bioanalysis method development, validation, and sample analysis: points to consider when conducting nonclinical and clinical studies in accordance with current regulatory guidances. J. Anal. Bioanal. Tech. 4.

Whitney SM, Andrews TJ. 1998. The CO2/O2 specificity of single-subunit ribulose-bisphosphate carboxylase from the dinoflagellate, Amphidinium carterae. Aust. J. Plant Physiol. 25: 131–138.

Whitney SM, Birch R, Kelso C, Beck JL, Kapralov MV. 2015. Improving recombinant Rubisco biogenesis, plant photosynthesis and growth by coexpressing its ancillary RAF1 chaperone. Proc. Natl. Acad. Sci. 112: 3564–3569.

Whitney SM, Houtz RL, Alonso H. 2011. Advancing our understanding and capacity to engineer nature’s CO2-sequestering enzyme, Rubisco. Plant Physiol. 155: 27–35.

Wilson RH, Hayer-Hartl M, Bracher A. 2019. Crystal structure of phosphoribulokinase from Synechococcus sp. strain PCC 6301. Acta Cryst. 75: 278–289.

Wilson RH, Martin-Avila E, Conlan C, Whitney SM. 2018. An improved Escherichia coli screen for Rubisco identifies a protein–protein interface that can enhance CO2-fixation kinetics. J. Biol. Chem. 293: 18–27.

Wilson RH, Whitney SM 2017. Improving CO2 fixation by enhancing rubisco performance. In: Alcalde M ed. Directed Enzyme Evolution: Advances and Applications. Cham: Springer International Publishing, 101–126.

Yamori W, Suzuki K, Noguchi K, Nakai M, Terashima I. 2006. Effects of Rubisco kinetics and Rubisco activation state on the temperature dependence of the photosynthetic rate in spinach leaves from contrasting growth temperatures. Plant Cell Environ. 29: 1659–1670.

Young JN, Heureux AMC, Sharwood RE, Rickaby REM, Morel FMM, Whitney SM. 2016. Large variation in the Rubisco kinetics of diatoms reveals diversity among their carbon-concentrating mechanisms. J. Exp. Bot. 67: 3445–3456.

Zhou W, Yang S, Wang PG. 2017. Matrix effects and application of matrix effect factor. Bioanalysis 9: 1839–1844.

Zhou Y, Gunn LH, Birch R, Andersson I, Whitney SM. 2023. Grafting Rhodobacter sphaeroides with red algae Rubisco to accelerate catalysis and plant growth. Nat. Plants 9: 978–986.

Zhou Y, Whitney S. 2019. Directed evolution of an improved Rubisco; in vitro analyses to decipher fact from fiction. Int. J. Mol. Sci. 20: 5019.

