## Supplemental Information S1-S4 for "RubiSpec-MS: Determination of Rubisco CO_2_/O_2_ Specificity Factor using Liquid Chromatography-Mass Spectrometry"

| **PAGE** | **CONTENTS** |
| --- | --- |
| S1 | Table of Contents |
| S2–S5 | Supplementary Tables |
| S6 | Supplementary References |

**Other supplementary materials for this manuscript include the following:**

**Tables S5** and **S6** (as .xls file)

**Table S1.** Plasmids used in this study.

| **Plasmid** | **Vector** | **Gene** | **GenBank ID** | **Source** |
| --- | --- | --- | --- | --- |
| pHue-PRK | pHue | *PRK* | ATN39931.1 | This work |
| pHue-6PGD | pHue | *6PGD* | NP_012053.3 | This work |
| pHue-RbcM | pHue | *rbcM* | WP_200290014.1 | This work |
| pTrc-*Syn*LS | pTrc | *Syn-rbcL* | WP_011242444.1 | (Mueller-Cajar & Whitney, 2008) |
|  |  | *Syn-rbcS* | WP_011242443.1 |  |
| pTrc-*Cs*LS | pTrc | *Cs-rbcLS* | KM464722.1 | This work |
| pCDF-*Nt*Assmbl | pCDF | *Nt-raf1* | XP_009764918.1 | (Buck *et al.*, 2022) |
|  |  | *Nt-raf2* | XP_009599433.1 |  |
|  |  | *Nt-rbcX* | XP_016509860.1 |  |
|  |  | *Nt-bsd2* | AYR18863.1 |  |
|  |  | *Nt-cpn60α* | XP_16509320.1 |  |
|  |  | *Nt-cpn60β* | XP_016490465.1 |  |
|  |  | *Nt-cpn20* | XP_0.16432480.1 |  |
| pET16-*Nt*LS | pET16 | *Nt-RbcL* | NP_054507.1 | This work |
|  |  | *Nt-RbcS* | 1EJ7_S |  |
| pACYC-*Syn*-^His6^Ubq-RbcS | pACYC | *Syn-RbcS* | WP_011242443.1 | (Mueller-Cajar & Whitney, 2008) |
| pACYC-*Cs* ^His6^Ubq-RbcS | pACYC | *Cs-RbcS* | WP_002721826.1 | This work |
| pACYC-*Nt* ^His6^Ubq-RbcS | pACYC | *Nt-RbcS* | 1EJ7_S | This work |
| pUSP2_ ^His6^USP2 | pUSP2 | *usp2* | NP_932759.1 | (Baker *et al.*, 2005) |

**Table S2.** Purified proteins concentrations. Monomer refers to single functioning unit of Rubisco (one active site).

| **Protein** | **ԑ (M^-1^ cm^-1^)** | **Molecular weight (g/mol)** | **A_280_** | **Monomer Concentration (μM)** |
| --- | --- | --- | --- | --- |
| *Rhodospirillum rubrum* Rubisco | 58330 | 50504 | 21.2 | 362.70 |
| *Nicotiana tabacum* Rubisco | 113345 | 67644 | 2.6 | 22.94 |
| *Synechococcus elongatus* PCC 6301 Rubisco | 90925 | 65737 | 22.6 | 248.56 |
| *Cereibacter sphaeroides* Rubisco | 107260 | 68457 | 2.1 | 19.58 |
| *Arabidopsis thaliana* Rubisco | 104000 | 67752 | 4.2 | 40.66 |
| *Spinacea oleracea* Rubisco | 106230 | 67022 | 3.5 | 32.95 |
| Phosphoribulokinase (PRK) | 36330 | 37615 | 16.9 | 465.18 |
| Phosphogluconate dehydrogenase (6PGD) | 69330 | 56081 | 22.2 | 320.21 |

**Table S3.** Matrix effect calculations. Matrix effect was calculated by comparing the peak area of an analyte in neat solvents and the peak area of the same analyte spiked into a simulated matrix at the same concentration (250 µM), *n* = 5.

|  | **Replicate** | **2PG peak area** | **3PGA peak area** | **RuBP peak area** |
| --- | --- | --- | --- | --- |
| **Analyte (250 μM) in neat solvents (A)** | 1 | 604933.938 | 64456.738 | 58669.008 |
|  | 2 | 611550.688 | 74314.555 | 69371.547 |
|  | 3 | 634290.625 | 79293.500 | 76406.719 |
|  | 4 | 679906.938 | 87233.758 | 79662.297 |
|  | 5 | 738920.938 | 92285.078 | 78073.125 |
| **Analyte spiked to 250 μM in sample matrix (B)** | 1 | 648803.115 | 80042.588 | 74414.771 |
|  | 2 | 655243.709 | 87447.466 | 79504.875 |
|  | 3 | 519903.646 | 80908.971 | 76678.420 |
|  | 4 | 541156.032 | 82775.333 | 78989.315 |
|  | 5 | 579659.861 | 87884.091 | 82628.435 |
| **Matrix Effect** | 1 | -7.252 | -24.180 | -26.838 |
|  | 2 | -7.145 | -17.672 | -14.607 |
|  | 3 | 18.034 | -2.037 | -0.356 |
|  | 4 | 20.407 | 5.111 | 0.845 |
|  | 5 | 21.553 | 4.769 | -5.835 |
| **Average Matrix Effect (%)** | | **9.12** | **-6.80** | **-9.36** |

**Table S4.** Raw chromatogram data for intraday precision calculations. Intraday precision was calculated from injecting the same standard mix (250 µM) 15 × on the same day. Relative standard deviation (RSD) % = (SD × 100)/mean.

|  | **Peak Area** | | |
| --- | --- | --- | --- |
| **Technical Replicate** | **2PG** | **3PGA** | **RuBP** |
| 1 | 537641.438 | 77458.945 | 59539.086 |
| 2 | 585300.750 | 85794.438 | 64526.281 |
| 3 | 616859.063 | 88049.320 | 68844.641 |
| 4 | 594241.813 | 88742.656 | 69180.867 |
| 5 | 641870.938 | 89317.539 | 69064.828 |
| 6 | 678023.313 | 85343.813 | 57522.230 |
| 7 | 665282.375 | 89042.063 | 64193.895 |
| 8 | 499152.969 | 81118.953 | 61342.508 |
| 9 | 455425.219 | 82431.711 | 55142.418 |
| 10 | 472617.813 | 83704.484 | 63894.621 |
| 11 | 484255.281 | 76338.570 | 56416.805 |
| 12 | 514481.469 | 85233.000 | 62851.469 |
| 13 | 524862.125 | 88676.648 | 57807.570 |
| 14 | 529672.375 | 90709.117 | 60812.582 |
| 15 | 549182.688 | 94008.391 | 69237.875 |
| **Intraday Precision (RSD%)** | **12.65** | **5.69** | **7.78** |

**Table S5.** $S_{C/O}$ determination of samples measured with the RubiSpec-MS method (see Excel file).

**Table S6**. $S_{C/O}$ determination of samples measured with the Radiolabeled RuBP method (see Excel file).

**Supplementary References**

**Baker RT, Catanzariti AM, Karunasekara Y, Soboleva TA, Sharwood R, Whitney S, Board PG 2005.** Using Deubiquitylating Enzymes as Research Tools. *Methods in Enzymology*: Academic Press, 540-554.

**Buck S, Rhodes T, Gionfriddo M, Skinner T, Yuan D, Birch R, Kapralov MV, Whitney SM. 2022.** *Escherichia coli* expressing chloroplast chaperones as a proxy to test heterologous Rubisco production in leaves. *J. Exp. Bot.* **74**: 664-676.

**Mueller-Cajar O, Whitney SM. 2008.** Evolving improved *Synechococcus* Rubisco functional expression in *Escherichia coli*. *Biochem. J.* **414**: 205-214.
